# Mutual Information Analysis of Neural Representations of Speech in Noise in the Aging Midbrain

**DOI:** 10.1101/619528

**Authors:** Peng Zan, Alessandro Presacco, Samira Anderson, Jonathan Z. Simon

## Abstract

Younger adults with normal hearing can typically understand speech in the presence of a competing speaker without much effort, but this ability to understand speech in challenging conditions deteriorates with age. Older adults, even with clinically normal hearing, often have problems understanding speech in noise. Earlier auditory studies using the frequency-following response (FFR), primarily believed generated by the midbrain, have demonstrated age-related neural deficits when analyzed using traditional measures. Here we use a mutual information paradigm to analyze the FFR to speech (masked by a competing speech signal) by estimating the amount of stimulus information contained in the FFR. Our results show, first, a broad-band informational loss associated with aging for both FFR amplitude and phase. Second, this age-related loss of information is more severe in higher frequency FFR bands (several hundred Hz). Third, the mutual information between the FFR and the stimulus decreases as noise level increases for both age groups. Fourth, older adults benefit neurally, i.e., show a reduction in loss of information, when the speech masker is changed from meaningful (talker speaking a language that they can comprehend, such as English) to meaningless (talker speaking a language that they cannot comprehend, such as Dutch). This benefit is not seen in younger listeners, which suggests age-related informational loss may be more severe when the speech masker is meaningful than when it is meaningless. In summary, as a method, mutual information analysis can unveil new results that traditional measures may not have enough statistical power to assess.

**New & Noteworthy:** Older adults, even with clinically normal hearing, often have problems understanding speech in noise. Auditory studies using the frequency-following response (FFR) have demonstrated age-related neural deficits using traditional methods. Here we use a mutual information paradigm to analyze the FFR to speech masked by competing speech. Results confirm those using traditional analysis, but additionally show that older adults benefit neurally when the masker changes from a language that they comprehend to a language they cannot.

## Introduction

Understanding speech in the presence of background noise becomes more challenging as humans age. Older listeners often report problems in listening to speech in noise even with clinically normal hearing sensitivity (Burke and Shafto 2008; Helfer and Freyman 2008). Behavioral studies have revealed age-related temporal processing deficits in a number of auditory tasks, such as pitch discrimination (Fitzgibbons and Gordon-Salantt 1996), gap-in-noise detection (Fitzgibbons and Gordon-Salant 2001) and recognition of speech in noise (Frisina and Frisina 1997; Gordon-Salant et al. 2006; He et al. 2008; Schneider and Hamstra 1999). These results suggest a temporal processing degradation in the auditory pathway, consistent with observed age-related changes in response latency and strength in midbrain (Anderson et al. 2012; Burkard and Sims 2002; Clinard and Tremblay 2013) and cortical evoked responses (Lister et al., 2011; Presacco et al., 2016a, 2016b).

The neural mechanism underlying age-related temporal auditory process deficits has also been investigated in animal studies: decreased release of inhibitory neurotransmitters, such as gamma-aminobutyric acid (GABA), in dorsal cochlear nucleus (Caspary et al. 2005; Parthasarathy and Bartlett 2011; Schatteman et al. 2008; Wang et al. 2009), inferior colliculus (IC) (Caspary et al. 1995) and auditory cortex (Juarez-Salinas et al. 2010; de Villers-Sidani et al. 2010) have been found in aging mammals. Because the spectro-temporal fine structure of speech is encoded by synchronous neural firing in midbrain, and the accurate processing of rapid fluctuations depends partly on inhibitory mechanisms, the representation of speech there also may deteriorate as a result of greater variability of neural firing (Walton et al. 1998; Yang et al. 1992) or loss of inhibition (Caspary et al. 2005, 2006; Walton et al. 1998). The midbrain frequency-following response (FFR), which tracks periodic components of speech or other sounds, may be detrimentally affected by the resulting neural jitter. In older listeners, jitter may be more prevalent than in younger listeners, as reflected by a decreased inter-trial response consistency (Anderson et al. 2012), or, as we hypothesize here, by increased entropy and decreased mutual information as defined in the context of information theory (Cover and Thomas 1991; Shannon 1948).

Mutual information, in particular, can be interpreted as a reduction in auditory response variability due to the presentation of a stimulus (Nelken and Chechik, 2007). It has been used to estimate transmission rates in the low-frequency fibers of the auditory periphery in bullfrog (Rieke et al. 1995), and applied to magnetoencephalography (MEG) auditory responses to continuous speech (Cogan and Poeppel 2011). Auditory information transmitted from midbrain to auditory cortex has been observed to show greater redundancy in older listeners compared to younger listeners (Bidelman et al. 2014). However, given that older listeners have a weaker midbrain response than younger listeners (Presacco et al. 2016a, 2016b), it remains an open question whether the aging midbrain itself processes more information or less information than younger listeners.

The current study is a mutual informational analysis of auditory midbrain FFR. A more traditional analysis (evoked response) of this dataset has already been published (Presacco et al., 2016a, 2016b). The goals of this new analysis are: 1) to describe these new and innovative methods in detail, 2) to demonstrate rich examples of their use, and 3) to demonstrate that the results are quite often stronger in statistical power than the more traditional methods. First it is shown that the new analysis replicates the most basic earlier findings, that older listeners’ midbrain FFR responses contain less auditory signal information about speech stimuli than younger listeners’, at the fundamental frequency (F_0_) of the FFR. We then generalize the analysis to harmonic frequencies, showing that speech information contained there is similarly degraded with age (and falls off more quickly in frequency), consistent with earlier findings (Anderson et al. 2012). Finally, we also show that when the speech stimuli are degraded by the addition of a competing talker, the stimulus information contained in the midbrain FFR is more sensitive to informational masking (competing speech in a familiar vs. unfamiliar language) in older listeners than in younger listeners.

## Materials and methods

### Subjects

The dataset used in this study has previously been described (Presacco et al., 2016a, 2016b). Seventeen younger listeners (3 men) between 18 and 27 years old (mean ± SD: 22.23 ± 2.27) and fifteen older listeners (5 men) between 61 and 73 years old (mean ± SD: 65.06 ± 2.30), recruited from the Maryland, Washington D.C. and Virginia areas, participated in the experiment. All subjects had clinically-normal hearing with air-conduction thresholds no greater than 25 dB hearing level (HL) from 125 to 4,000 Hz bilaterally and no interaural asymmetry. All of them were native English speakers and were free of neurological or middle-ear disorders, and none of them spoke or understood the Dutch language. All participants were paid for their participation, and each of them gave written informed consent before the experiment. The experimental protocol and all procedures were reviewed and approved by the Institutional Review Board of the University of Maryland.

### Stimuli and EEG recording

The stimulus was a single speech syllable, a 170-ms /da/ (Anderson et al. 2012), synthesized at a 20-kHz sampling rate with a Klatt-based synthesizer (Klatt 1980) with a 100-Hz F_0_. The syllable was chosen because it comprises both transient and steady-state components, the stop consonant /d/ is rich in phonetic information, and its perception is sensitive to background noise (Miller and Nicely 1955). Its waveform and spectrum are shown in Figure 1. The speech syllable was presented diotically at 75 dB SPL with a repetition rate of 4 Hz. Stimuli were presented with alternating polarities to allow cancellation of potential stimulus artifact by summing the responses to each pair (Aiken and Picton 2008). The stimulus was presented to subjects both in quiet and in noise. For the noise conditions, a story narrated by a female competing speaker in either English or Dutch was used as a masker (a 1-minute duration segment, continuously looped). The English story was an excerpt from A Christmas Carol by Charles Dickens (http://www.audiobooktreasury.com/a-christmas-carol-by-charles-dickens-free-audio-book/), and the Dutch story was Aljaska en de Canada-spoorweg by Anonymous (http://www.loyalbooks.com/book/Aljaska-en-de-Canada-spoorweg). For each of the two masker types, four signal-to-noise ratio (SNR) levels, +3, 0, −3, and −6 dB SNR, were created by using the logarithm of the ratio between root-mean-squared values of syllable /da/ and the long-duration masking speech. All stimuli were presented by insert earphones (ER1, Etymotic Research, Elk Grove Village, IL) via Xonar Essence One (ASUS, Taipei, Taiwan) using Presentation software (Neurobehavioral Systems, Berkeley, CA). FFRs were recorded at a sampling frequency of 16,384 Hz using the ActiABR-200 acquisition system (BioSemi B.V., Amsterdam, Netherlands) with a standard vertical montage of five electrodes (Cz active, forehead ground common mode sense/driven right leg electrodes, earlobe references), and the recorded signal was filtered online by a band-pass filter with a cutoff band of 100 Hz to 3,000 Hz. During the 2-hour recording session, subjects sat in a recliner and watched a silent captioned movie of their choice to facilitate a relaxed but wakeful state. For each of the nine conditions (1 quiet + 2 masker languages × 4 SNRs), at least 2,300 trials of response (to repetitions of syllable /da/) were recorded.

**Figure 1.**
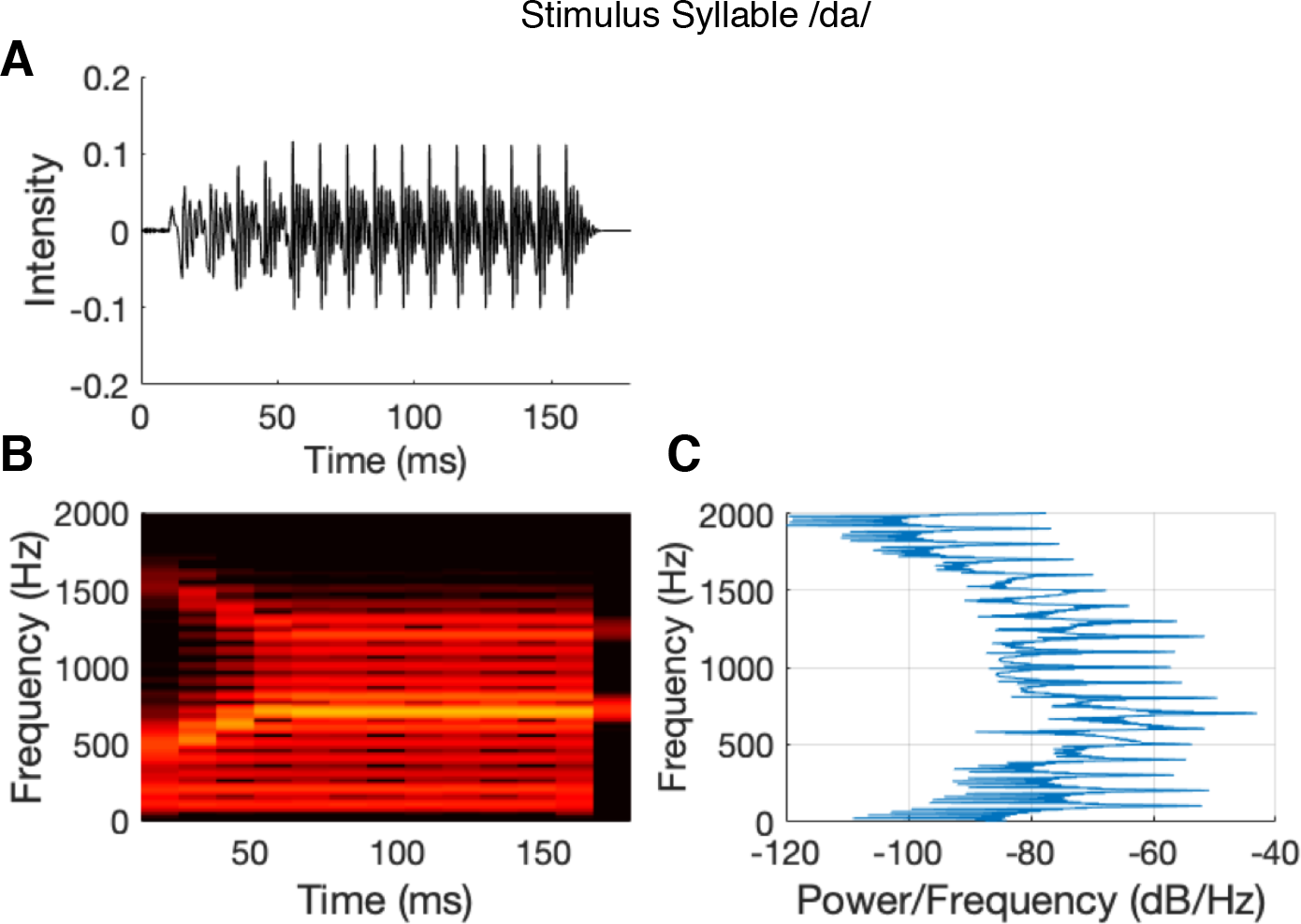
A: Stimulus waveform, B: spectrogram, and C: power spectral density of 170-ms syllable /da/. The locations of the horizontal peaks in C indicate that the syllable has a fundamental frequency of 100 Hz with harmonic peaks at its multiples (Anderson et al. 2012, 2013)

### Data analysis

#### Encoding response amplitude

The EEG recordings were first converted into MATLAB format with the function *pop_biosig* from EEGLab (Delorme and Makeig 2004), and all remaining analyses performed in MATLAB (version 2017b; Mathworks, Natick MA). The EEG recordings were band-pass filtered offline, to remove low-frequency neural oscillations, from 70 Hz to 2000 Hz with a linear-phase FIR filter with low-pass transition band of 65-70 Hz and high-pass transition of 2000-2100 Hz. Filter delays were compensated by processing the data in both forward and backward directions, using the Matlab function *filtfilt* (Mathworks, Natick MA). The response of each trial was analyzed in the time window −47 ms to 170 ms with respect to stimulus onset. Within this window, the response of each trial was band-pass filtered with linear-phase FIR filters of order 200, designed using least-square error minimization, into frequency bands centered at harmonics of 100 Hz, i.e., 100, 200, …, 600 Hz, to investigate the midbrain representations of harmonics. Harmonics at or above 700 Hz, the first formant of the steady-state portion of the stimulus, were excluded from analysis. Sweeps with amplitudes larger than ±30 μV were excluded, allowing 2000 artifact-free sweeps to be used. To eliminate any possible electrical feedthrough artifacts, a 10-ms temporal response function centered at 0 ms with reference to the stimulus onset time was estimated per trial, and its contribution was subtracted from the response (Maddox and Lee 2018). Additionally, since two consecutive sweeps were always presented with opposite polarities, their responses were averaged into one effective sweep, leading to 1000 such pair-averaged sweeps per subject and per condition that were then used for the analysis; the results for the same sweeps, with artifacts removed but not averaged (2000 per condition) are presented in the Appendix. For each of the two analysis regions, the response waveforms were extracted from each sweep for every subject, for each of the nine conditions and 6 frequency bands.

Under each condition, for each subject and frequency band, a response matrix was obtained of size 1000 trials × *T* samples where *T* is the sample length of observation window. In addition to the entire response window 0-170 ms, the responses were also partitioned into two regions based on the acoustic properties of the syllable /da/, i.e., the transition (15-65 ms) and steady-state (64-170 ms) for analysis of masker type influence on the response at 100 Hz. Here *T* = 2853 samples for the entire response region, *T* = 804 samples for the transition region, and *T* = 2049 samples for the steady-state region. The response amplitudes at each sample were subdivided into *N* bins with the boundaries of the bins chosen so that approximately equal numbers of samples were assigned in each bin; each sample was then associated with its bin index (from 1 to *N*). The boundaries were chosen individually based on each subject’s response. Different values of *N* ∈ {4, 8, 16, 32, 64, 128} were evaluated to verify a lack of any interaction with age (*F*_(5,180)_ = 0.46, *p* = 0.809 and *F*_(5,180)_ = 0.18, *p* = 0.970 by ANOVA test on interaction of *age* × *bin number* for amplitude and phase information, respectively). A final choice of *N* = 32 bins was selected as an optimal trade-off between increased resolution between bins and decreased samples per bin due to limited samples (too few bins or too few samples per bin both lead to estimation bias). The choice of 32 bins gave more than 30 samples/bin, on average, to estimate the conditional probability distribution.

#### Encoding response phase

For every sweep in each region, the phase for each frequency band was computed by first applying the Hilbert transform to the band-passed signal and then computing the phase of the resultant complex (analytic) signal, i.e.,

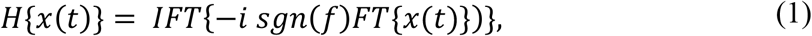

where *FT* is the Fourier Transform, *f* is the frequency basis of the Fourier Transform, *sgn*(*f*) is the algebraic sign of *f*, and *IFT* is the inverse Fourier Transform. Then

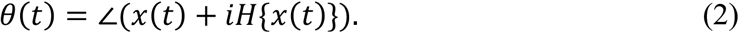

The phase-locking value (PLV) of the response in any single band can be computed as

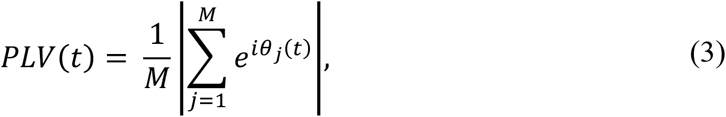

where *θ*_*j*_(*t*) is the phase of *j*^*th*^ trial at sample time *t*, and *M* is the number of trials.

The set of phase responses *θ*_*j*_(*t*) obtained for each frequency band, were also subdivided in to *N* = 32 bins, analogously to encoding the amplitude response; here the phase samples were divided into bins of width 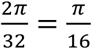 with each sample encoded by its bin index (from 1 to *N*).

#### Mutual information

Under each condition, for each subject and each frequency band, the mutual information between stimulus and amplitude, and mutual information between stimulus and phase were estimated based on those integer-encoded responses. The response probability distribution was estimated as above (bin index for each of the *T* samples over 1000 trials). The conditional distribution of *P*(*Y*|*X*) was drawn from response samples at the same latency from 1000 trials. The mutual information can then be estimated by the entropy of the response, whether amplitude or phase, minus the conditional entropy of the response given the (uniformly distributed) stimulus:

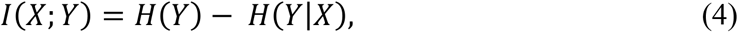

where *X* represents the stimulus distribution, and *Y* is the response distribution, whether amplitude or phase. *H*(*Y*) is the entropy of the response,

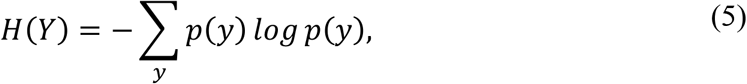

where *p*(*y*) is the probability of observing the response value *y*. *H*(*Y*|*X*) is the entropy of the response conditioned by the stimulus *X* and is given by:

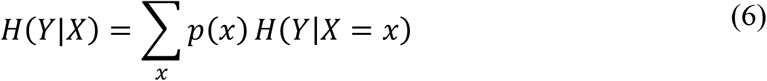

where

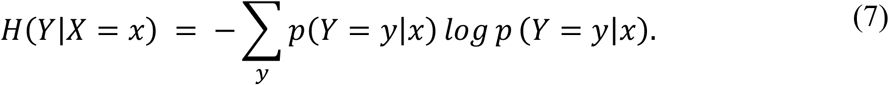

The stimulus *X* is the amplitude or phase at each time point. The probability distribution of *x* is unknown, but here assumed to be uniform (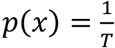, a constant, so each bin contains roughly the same number of stimulus value instances) for two reasons. First, when the actual stimulus distribution is unknown, this assumption minimizes estimation bias (Nelken and Chechik 2007). Second, while there is not yet evidence for any particular distribution (e.g., Gaussian or Laplacian), the assumption of uniform distribution was employed for stimulus amplitude by Cogan and Poeppel (2011) with encouraging results. Then, equation (3) becomes

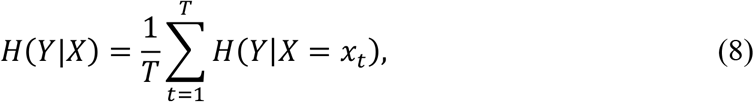

where *x*_*t*_ is the amplitude or phase bin at sample *t*.

To illustrate, consider an analysis of the quiet condition over the steady-state region, which encompasses the time window from 64 ms to 189 ms with respect to stimulus onset, i.e., 2,049 samples, giving *T* = 2049 and 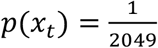 for every value of *t*.

The distribution of the response, *P*(*Y*), is estimated for each subject with all bin-index-encoded samples in each of the 1,000 trials. The conditional distribution of *Y* given *x*_*t*_, *P*(*Y*|*x*_*t*_), is estimated with 1,000 samples from trials at time point *t*. Then the conditional entropy is given by

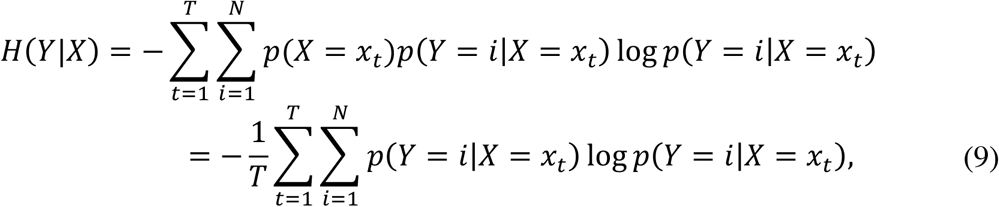

where *i* ∈ {1, 2, …, *N*} is the bin number and *N* is the number of bins. The mutual information is therefore,

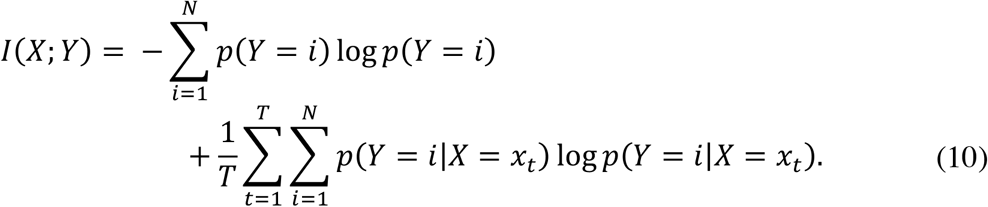

#### Statistics

To examine the effects of aging, frequency, masker type and SNR level, multiple *t-*tests with correction were performed, separately for both amplitude and phase information. To facilitate analyzing the information at fundamental frequency, linear models were constructed to test effects from interactions between aging and other factors, namely masker type and SNR level, with mathematical form *I* ~ *age* × *masker type* + *age* × *SNR*. Tests were performed for both amplitude and phase, and for different temporal regions, separately. To test masker type influence within group, the mutual information difference between Dutch and English maskers for each subject was modeled as *I*_*Dutch*_ – *I*_*English*_ ~ *SNR*, and the positivity of intercept was tested for both amplitude and phase, and for different temporal regions, separately. The results were justified by *t*-tests on the intercept of linearly fitted regression lines for each subject, and similar analysis for PLV.

Linear models with only fixed effects were analyzed in R (R. Core Team 2017) using the function *lm*, which reports the model significance using an *F*-test on the constructed model vs. the null model with only the intercept, and the significance of influence from fixed-effect factors with separate *t*-tests on the slope of each factor. The assumption of Homoscedasticity of the linear models was examined by global validation of linear model assumptions using toolbox *gvlma* (Peña and Slate 2006) in R. Responses at harmonic frequencies were analyzed using *t*-tests. False discovery rate correction (FDR) (Benjamini and Hochberg 1995), to correct for multiple comparisons, was applied when appropriate.

Where appropriate, *t*-tests for significance are supplemented with effect size (Cohen’s *d*) and its 95% confidence interval (CI). When the CI excludes zero, this is alternate evidence that the result is statistically significant (i.e., the effect size is significantly greater than zero at an α level of 0.05). Note, however, that the effect size analysis is not compensated for multiple comparisons even when the *p*-value is.

The effective high-frequency cutoff for any frequency-decreasing statistical measure is defined to be the frequency at which the measure is not significantly higher than the noise floor (pure estimation bias). The noise floor is estimated using the same mutual information method as used elsewhere, but instead using responses to quiet intervals between stimuli.

## Results

Here, we report results from the mutual information analysis of pair-averaged-polarity responses; the analogous analysis based on single sweeps is reported in the Appendix. Because our algorithm takes into account variations across trials, pair-averaging provides less variation and thus higher mutual information. Except for this overall scaling of mutual information, the results are typically comparable.

### Information in FFR amplitude

#### Amplitude information at 100 Hz

For the amplitude response at 100 Hz, to examine masker type and SNR interactions with both age groups, the linear model, *I* ~ *age* × *masker type* + *age* × *SNR* is tested. It is significant (*F*_5,250_ = 4.99, *p* < 0.001 for the entire region; *F*_5,248_ = 2.93, *p* = 0.014 for the transition region; *F*_5,249_ = 6.11, *p* < 0.001 for the steady-state region). Outliers that would otherwise cause the homoscedasticity requirement to be violated are excluded (2 samples from the transition region and 1 sample from the steady-state region, respectively). Results show no significant interactions between *age* and *masker type* (*t*_(250)_ = 0.53, *p* = 0.587 for the entire region; *t*_(248)_ = 0.15, *p* = 0.884 for the transition region; *t*_(249)_, *p* = 0.773 for the steady-state region), or between *age* and *SNR* (*t*_(250)_ = 0.79, *p* = 0.428 for the entire region; *t*_(248)_ = 0.46, *p* = 0.645 for the transition region; *t*_(249)_ = 0.87, *p* = 0.386 for the steady-state region). A linear model with no interactions was then constructed and tested, i.e., *I* ~ *age* + *masker type* + *SNR*. The model itself is significant (*F*_3,252_ = 8.05, *p* < 0.001, *F*_3,250_ = 4.84, *p* = 0.003, *F*_3,251_ = 9.96, *p* < 0.001 for the entire region, the transition and steady-state regions, respectively). Comparisons between the models show that younger listeners’ responses contain significantly more information than older listeners’ responses in the whole and steady-state regions (*t*_(252)_ = 4.24, *p* < 0.001 and *t*_(251)_ = 4.99, *p* < 0.001 respectively), and that information increases as SNR increases (*t*_(252)_ = 2.37, *p* = 0.018 for the entire region; *t*_(250)_ = 2.86, *p* = 0.005 for the transition region; *t*_(251)_ = 2.15, *p* = 0.033 for the steady-state region).

Since the stimulus has a fundamental frequency of 100 Hz and the phase-locking of FFR is more robust in low frequencies than in high frequencies (Zhu et al. 2013), the 100-Hz FFR may contain significantly more information than its harmonics. To rule out the possibility that significant contributions to mutual information derive from averaging the opposite polarities, the same mutual information analysis is performed on single trials, where similar results are observed (see Appendix). Figure 2A displays the mutual information as a function of SNR level. Older listeners not only have a noticeably lower amount of information than younger listeners, but also extract more speech information when the masker is Dutch than for English. To eliminate within-subject variance, a linear regression line of information-by-SNR is fitted for each subject and its y-intercept and slope were analyzed, with results illustrated in Figure 2. A one-tailed *t*-test (younger > older) on the y-intercept shows a significantly larger amount of information in younger than older listeners for the English masker (*t*_(30)_ = 1.71, *p* = 0.048; *d* = 0.75, 95% *CI* = [0.032, 1.469]). The difference is not significant for Dutch (*t*_(30)_ = 1.41, *p* = 0.102; *d* = 0.51, 95% *CI* = [−0.195, 1.216]) (but as will be seen below, it does become significant for higher harmonic frequencies). Both age groups demonstrate decreasing information with worsening SNR: a one-tailed *t*-test on the negativity of the regression slope shows information loss for all cases except for older listeners with the Dutch masker (*t*_(16)_ = 3.42, *p* = 0.002 and *t*_(16)_ = 2.54, *p* = 0.013 for younger listeners for English and Dutch maskers, respectively, and *t*_(14)_ = 2.32, *p* = 0.027; *d* = 0.60, 95% *CI* = [2.55×10^−5^, +∞] and *t*_(14)_ = 2.35, *p* = 0.059; *d* = 0.61, 95% *CI* = [1.92×10^−5^, +∞] for older listeners). No significant difference is seen between the slopes across age groups (one-tailed *t*-test: *t*_(30)_ = 1.28, *p* = 0.106; *d* = 0.55, 95% *CI* = [−0.155, 1.260] for the English masker; *t*_(30)_ = 1.20, *p* = 0.120; *d* = 0.73, 95% *CI* = [0.018, 1.452] for the Dutch masker, though the effect size CI is consistent with significance in the last case.

**Figure 2.**
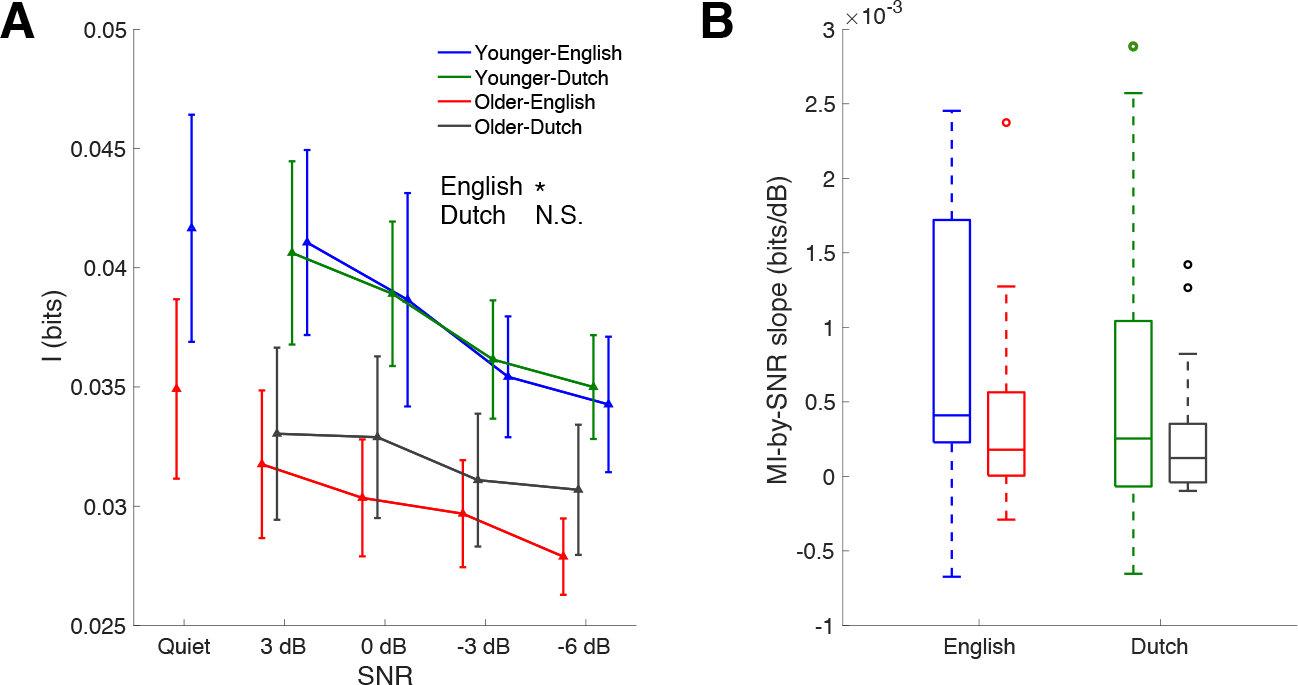
Mutual information between stimulus and response amplitude as a function of noise level for each age group and masker condition (masker language). A: Mutual information at the fundamental frequency as a function of noise level (quiet condition and 4 SNR levels) with blue and green for younger listeners (English and Dutch maskers, respectively), and red and gray for older listeners (English and Dutch maskers, respectively). The response in younger listeners conveys noticeably more information than the response in older listeners for the English masker condition, but the difference for Dutch is not significant at 100 Hz. Older listeners show consistently higher mutual information for the Dutch masker than for the English (the younger listeners show no consistent difference), but the difference is not significant at 100 Hz. B: The MI-by-SNR slopes of the previous plots show decreasing trends as SNR worsens, regardless of masker type, for both age groups. Younger listeners show a steeper decrease than older listeners but the difference is not significant at 100 Hz response. Error bars indicate one standard error of mean (SEM). (∗ *p* < 0.05)

#### Amplitude information in harmonics of 100 Hz

To analyze aging-associated informational loss for the harmonics (200 to 600 Hz), similar tests are performed on mutual information in responses of these frequencies (analysis stops before 700 Hz, which represents the first formant of the steady-state portion of the stimulus). In each harmonic, a linear regression line of mutual information as a function of SNR is fitted for each subject under each masker type. First the y-intercept of the fitted line at 3 dB is analyzed for group differences (see Figure 3).

**Figure 3.**
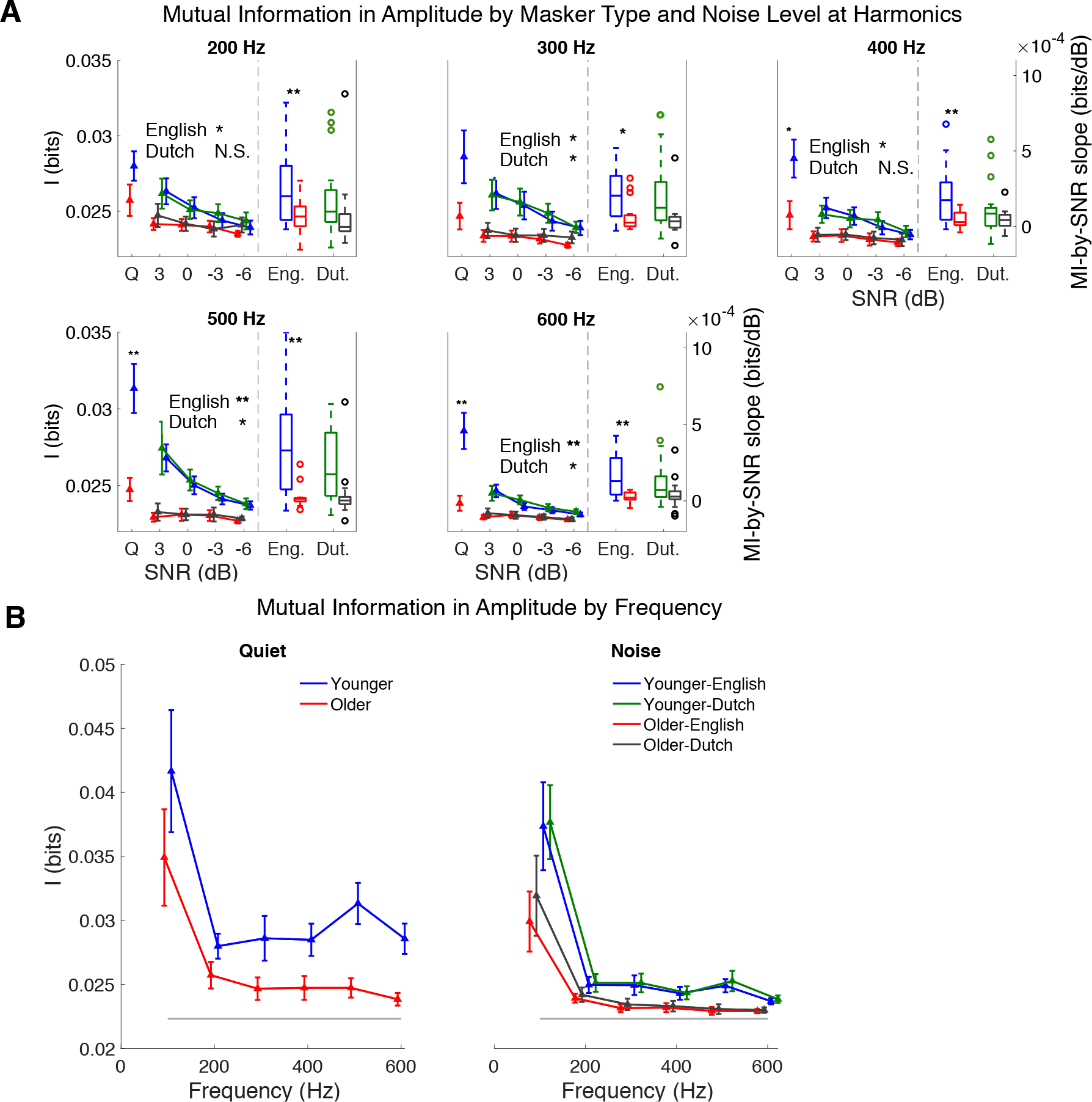
A: Mutual information for amplitude across frequency bands from 200 Hz to 600 Hz (separate subplot for each band). Within each subplot, the left panel shows the mutual information as a function of SNR, separately for age group and masker type. For the quiet condition, any asterisks above the error bars indicate the significance levels of group differences; text and any asterisks above the plots demonstrate significance levels of group differences in the corresponding masker types. Only younger listeners convey a significant amount of information in the higher harmonics. In the right panels, the bar plots depict the linearly fitted decreasing slopes (of the plots shown in the left panel) for the different age groups and masker types. In most bands, the mutual information decreases at a faster rate in younger listeners than in older. B: Overall, both in quiet (left) and averaged over SNR levels (right), mutual information decreases with increasing frequency (except for a single increase at 500 Hz for younger listeners). For older listeners, the decreasing trend in mutual information levels off at 300 Hz, which is lower than the frequency (>600 Hz) at which amplitude information levels off in younger listeners. The lower gray line represents the noise floor. Error bars indicate one SEM. (∗ < 0.05,∗∗ < 0.01)

One-tailed (younger > older) *t*-tests (with FDR correction) and effect size analysis on the y-intercept (corresponding to 3 dB SNR) of the line fit across all SNR levels suggest that the aging midbrain contains significantly less information than the younger midbrain in all frequencies from 100 to 600 Hz in the English masker condition. For *p*-values near 0.05 (see Table 1), effect size analysis is further applied. For the English masker condition, the 100-Hz condition shows consistent significance from both tests (*t*_(30)_ = 1.714, *p* = 0.048; *d* = 0.75, 95% *CI* = [0.032, 1.469]), and similarly for the Dutch masker condition at 300 Hz ( *t*_(30)_ = 2.05, *p* = 0.049; *d* = 1.236, 95% *CI* = [0.478, 1.993]), 500 Hz ( *t*_(30)_ = 2.27, *p* = 0.047; *d* = 0.787, 95% *CI* = [0.0663, 1.507]) and 600 Hz (*t*_(30)_ = 2.26, *p* = 0.047; *d* = 1.053, 95% *CI* = [0.312, 1.794]) (see also Figure 3A). In the English masker condition, one-tailed *t*-tests on fitted regression line slopes of younger listeners compared to older listeners show significantly steeper slopes for younger listeners compared to older listeners at frequencies from 200 to 600 Hz (all *p*-values are smaller than 0.05). All *p*-values of multiple comparisons are corrected. Overall, higher harmonics contain significant information only for younger listeners, and the difference in information between the two age groups becomes more statistically significant as the observed frequency increases, which is consistent with the linear model analysis, where age × frequency interaction is significant.

**Table 1.** Amplitude information: one-tailed *t*-test (younger > older) results applied to the fitted y-intercepts (3 dB values) and slopes from the linear regression analysis of mutual information (for response amplitude) as a function of SNR, for each harmonic. *p*-values are corrected for multiple comparisons by FDR correction. Boldfaced entries indicate the corresponding tests are statistically significant.

| Harmonic<br>(Hz) | Quiet<br>(Y>O) |  | English masker (Y>O) |  |  |  | Dutch masker (Y>O) |  |  |  |
| --- | --- | --- | --- | --- | --- | --- | --- | --- | --- | --- |
|  |  |  | y-intercept |  | slope |  | y-intercept |  | slope |  |
| | $t(30)$ | $p$ | $t(30)$ | $p$ | $t(30)$ | $p$ | $t(30)$ | $p$ | $t(30)$ | $p$ |
| 100 | 1.056 | 0.150 | **1.714** | **0.048** | 1.275 | 0.106 | 1.405 | 0.102 | 1.199 | 0.120 |
| 200 | 1.542 | 0.080 | **1.965** | **0.035** | **2.737** | **0.008** | 1.223 | 0.115 | 1.262 | 0.120 |
| 300 | 1.871 | 0.053 | **2.242** | **0.024** | **2.390** | **0.014** | **2.051** | **0.049** | 2.019 | 0.108 |
| 400 | **2.271** | **0.030** | **2.261** | **0.024** | **2.835** | **0.008** | 1.767 | 0.066 | 1.502 | 0.108 |
| 500 | **3.449** | **0.003** | **3.671** | **0.003** | **3.677** | **0.002** | **2.268** | **0.047** | 1.830 | 0.108 |
| 600 | **3.412** | **0.003** | **3.340** | **0.003** | **3.565** | **0.002** | **2.259** | **0.047** | 1.629 | 0.108 |

#### Amplitude information frequency limits

As seen in Figure 3B, the stimulus information contained in the response amplitude decreases with frequency for both age groups. The frequency-decreasing measure used here is the amplitude information’s y-intercept at 3dB of the fitted MI-by-SNR regression line. The frequency bands below 700 Hz are analyzed, separately for different masker types. The measure at 600 Hz for older listeners is not statistically distinguishable from the noise floor (*t*_(14)_ = 1.72, *p* = 0.107 by one-sample *t*-test). For younger listeners, the measure is significantly higher than the noise floor at all frequencies (*t*_(30)_ = 3.34, *p* = 0.002 for English masker; *t*_(30)_ = 2.26, *p* = 0.016 for Dutch masker (younger > older), both at 600 Hz where the lowest information is observed), i.e., the information for younger listeners has not yet reached floor by 600 Hz. In contrast, the cutoff frequency for older listeners is 300 Hz: the information measure at 300 Hz is not significantly greater than that at 600 Hz (*t*_(14)_ = 1.32, *p* = 0.130 under the English masker; *t*_(14)_ = 1.65, *p* = 0.095 under the Dutch masker). Therefore, results suggest a lower frequency limit of in amplitude information of 300 Hz for older listeners than that of beyond 600 Hz for younger listeners.

#### Effect of masker type on amplitude information

As seen in Figure 2B, older listeners demonstrate a slower fall-off in amplitude information as a function of SNR when the noise masker is Dutch than for English. To test for any potential amplitude information benefit from the Dutch masker over the English masker, the difference in information between the Dutch and English maskers is calculated for each subject in all SNR levels (for both transition and steady-state regions), and a linear model of *I*_*Dutch*_ – *I*_*English*_ ~ *SNR* shows a significantly positive intercept for older listeners in the transition region (*t*_(57)_ = 2.35, *p* < 0.001 with 2 samples omitted) but not in the steady-state region (*t*_(56)_ = 1.38, *p* = 0.173 with one sample omitted). Younger listeners, however, do not show a significant positive intercept in either transition (*t*_(65)_ = 1.90, *p* = 0.061 with one sample omitted) or steady-state region (*t*_(66)_ = −0.60, *p* = 0.549). Samples were omitted from the tests to satisfy the homoscedasticity requirement. A regression line was fitted as a function of SNR to reduce within-subject variance. Using a one-tailed *t*-test on the y-intercept (effective mutual information benefit at 3 dB SNR) of the regression line against zero, the mutual information benefit from the Dutch masker over the English masker is significantly higher for older listeners in the transition region (*t*_(14)_ = 2.35, *p* = 0.017), but not the steady-state region (*t*_(14)_ = 1.67, *p* = 0.058). No significant benefit is found for younger listeners in either region (*t*_(16)_ = 1.17, *p* = 0.130 and *t*_(16)_ = 0.51, *p* = 0.307 for transition and steady-state region, respectively). The regression slope is not significantly positive or negative for either group (*p* > 0.05 by two-tailed *t*-tests), as seen in the bar plots in the right panels of figure 4C and 4D.

**Figure 4.**
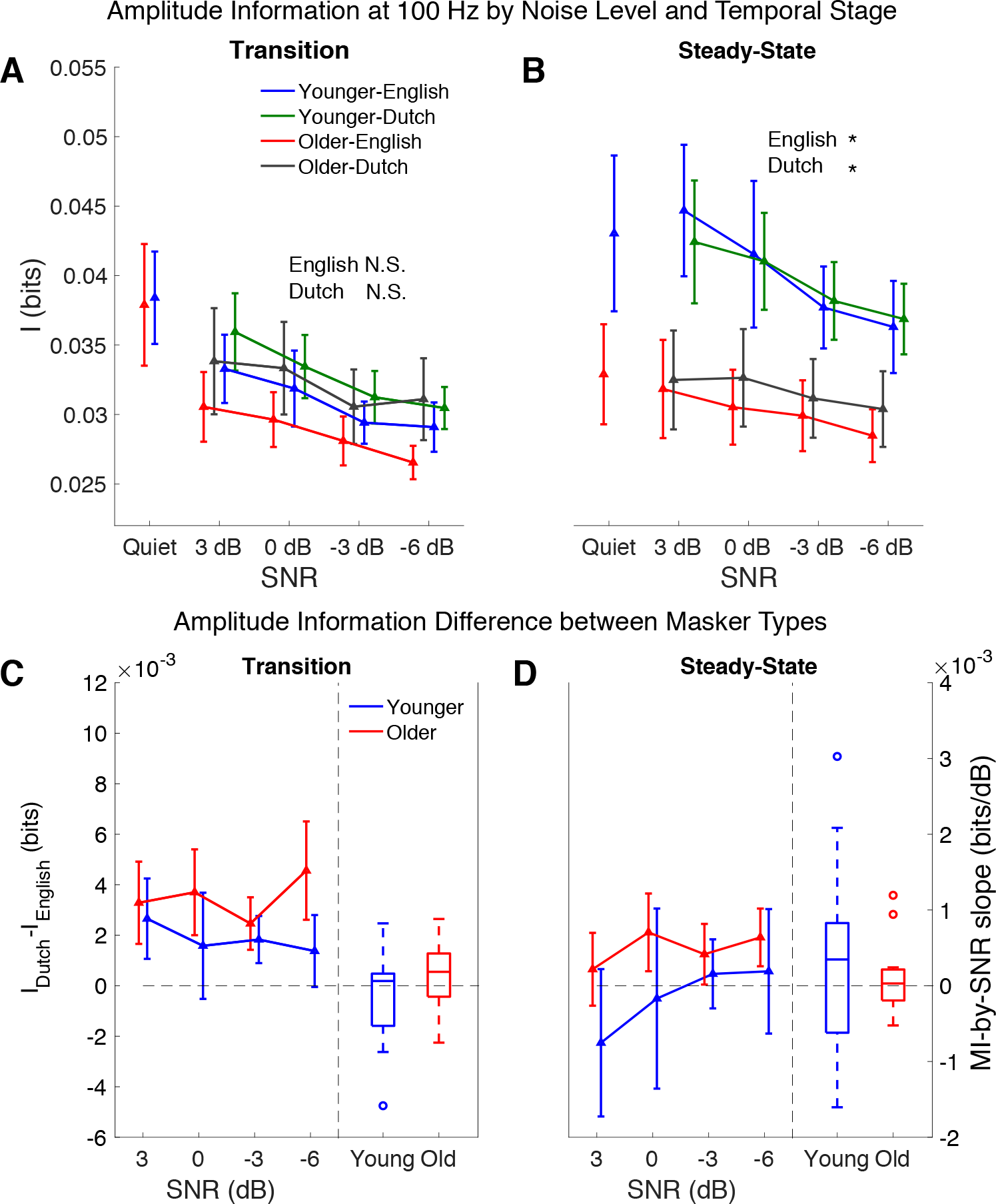
Mutual information of amplitude response by masker type and response region for younger listeners in blue (English) and green (Dutch) and older in red (English) and gray (Dutch). A and B demonstrate the mutual information as a function of SNR in the transition and steady-stage regions, respectively. In the steady-state region, group differences are significant for both masker types, indicated by asterisks. C and D illustrate the mutual information difference between masker types (denoted *I*_Dutch_ – *I*_*English*_) in the transition region and steady-state region, respectively. In each plot, the left panel displays information as a function of SNR, and the right panel displays a bar plot showing the slopes of the linear fits. The y-intercepts (corresponding to the fit at 3 dB SNR) are tested against 0 bits. Older listeners show significant benefit from the Dutch masker over English (denoted by asterisk), but only in the transition region. Error bars in all plots indicate SEM. (∗ *p* < 0.05)

### Phase-locking value

Phase-locking value (PLV) is a traditional measure of inter-trial coherence for a narrow-band response. Figure 5 shows the grand average of PLV at 100 Hz by age and masker condition. Older listeners have lower phase-locking values than the younger listeners (*t*_(30)_ = 2.62, *p* = 0.007 for one-tailed *t*-test) on the averaged phase-locking values across time and SNR levels. By one-tailed *t*-tests (*PLV*_*Dutch*_ – *PLV*_*English*_ > 0), older listeners have significantly higher PLV under Dutch masking than English (*t*_(14)_ = 2.74, *p* = 0.008 for transition region; *t*_(14)_ = 1.80, *p* = 0.047 for steady-state region), while younger listeners’ PLV is not significantly affected by informational masking (*t*_(16)_ = 1.67, *p* = 0.058 for transition region; *t*_(16)_ = 0.05, *p* = 0.479 for steady-state region).

**Figure 5.**
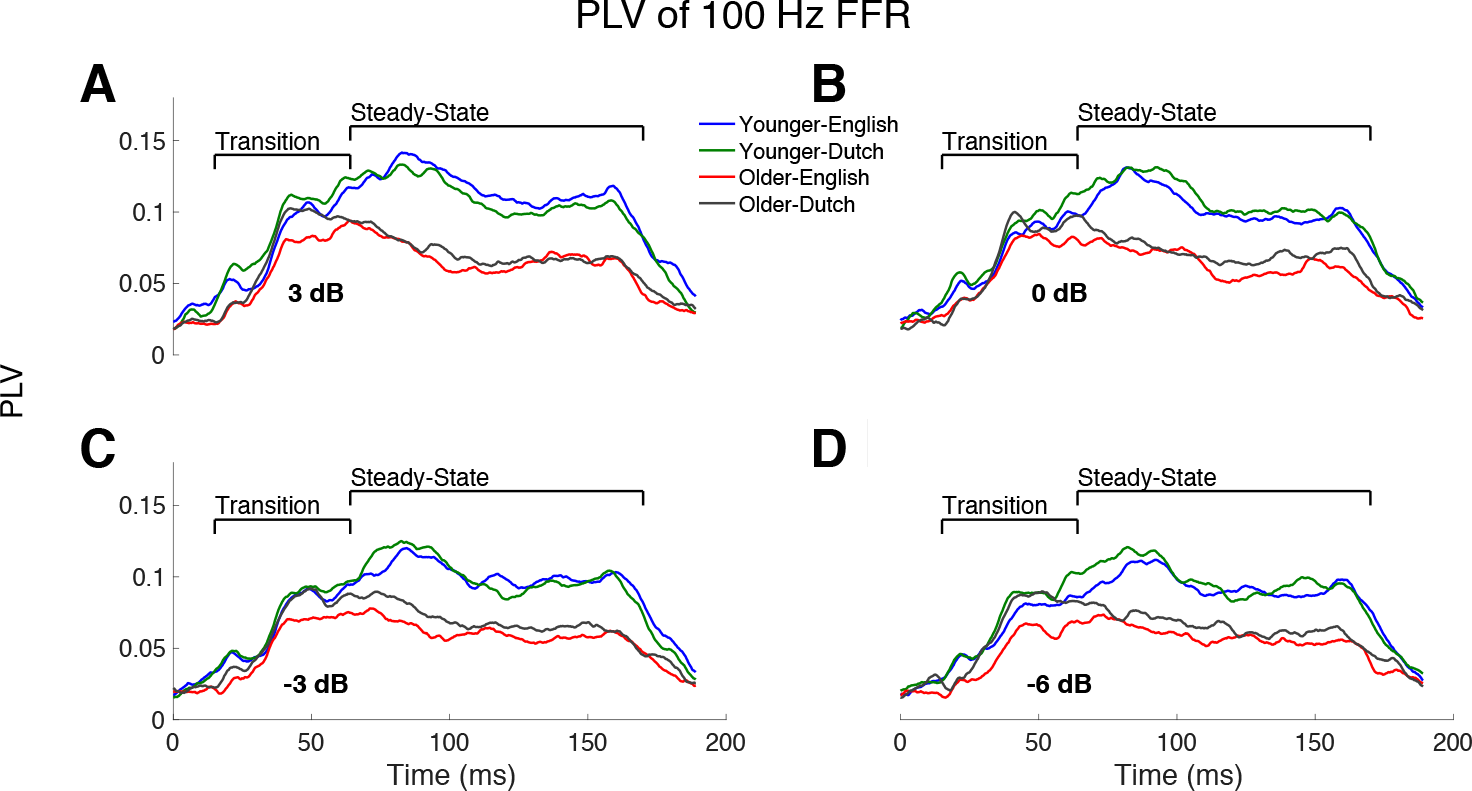
The PLV of the 100-Hz FFR is shown for all SNR levels, averaged across subjects, with colors indicating age and masker language, with younger listeners in blue (English) and green (Dutch) and older listeners in red (English) and gray (Dutch). A-D correspond to the four SNR levels: 3, 0, −3, −6 dB SNR. Younger listeners have visibly higher phase locking than older listeners. Older listeners have significantly better phase locking for the Dutch masker than for the English.

### Information in phase of FFR

#### Phase information at 100 Hz

For the phase response at 100 Hz, the linear model, *I* ~ *age* × *masker type* + *age* × *SNR* is significant (*F*_(5,250)_ = 5.45, *p* < 0.001 for the entire region; *F*_(5,248)_ = 3.27, *p* = 0.007 for the transition region; *F*_(5,248)_ = 6.24, *p* < 0.001 for the steady-state region). Outliers are excluded to satisfy homoscedasticity assumption (2 samples from transition and 2 samples from steady-state regions). Results show no significant interactions between *age* and *masker type* (*t*_(250)_ = 0.56, *p* = 0.578 for the entire region; *t*_(248)_ = 0.22, *p* = 0.825 for the transition region; *t*(248) = 0.06, *p* = 0.954 for the steady-state region), and between age and SNR (*t*_(250)_ = 0.86, *p* = 0.393 for the entire region; *t*_(248)_ = 1.05, *p* = 0.297 for the transition region; *t*_(248)_ = 0.66, *p* = 0.511 for the steady-state region). A linear model with no interactions was then constructed and tested, i.e., *I* ~ *age* + *masker type* + *SNR*. The model itself is significant (*F*_(3.252)_ = 8.77, *p* < 0.001, *F*_(3,250)_ = 5.08, *p* = 0.002, *F*(3.250) = 10.32, *p* < 0.001 for the entire region, the transition and steady-state regions, respectively). Comparisons show that younger listeners’ responses contain significantly more information than older listeners’ responses in the steady-state region (*t*_(252)_ = 4.52, *p* < 0.001 for the entire region; *t*_(250)_ = 2.12, *p* = 0.035 for the transition region; *t*_(250)_ = 5.19, *p* < 0.001 for the steady-state region), and that information increases as SNR increases (*t*_(252)_ = 2.31, *p* = 0.022 for the entire region; *t*_(250)_ = 2.63, *p* = 0.009 for the transition region).

Mutual information between stimulus and the response phase is analyzed analogously to that of the response amplitude. Phase information at 100 Hz is examined separately from the higher harmonics. To examine the effect of age and noise level, a linear regression line is fitted for information-by-SNR for each subject in both noise contents. The fitted y-intercept is compared for group differences. A one-tailed t-test (younger > older) effect size analysis on the y-intercept shows a significantly larger amount of information in younger than older listeners for the English masker (*t*_(30)_ = 1.80, *p* = 0.041; *d* = 0.82, 95% *CI* = [0.095, 1.540]); the difference is not significant for Dutch (*t*_(30)_ = 1.36, *p* = 0.092; *d* = 0.58, 95% *CI* = [−0.133, 1.284]) (Figure 6A). Both age groups demonstrate decreasing information with worsening SNR: a one-tailed t-test on the negativity of the regression slope shows information loss, however, the negativity is not significant for older listeners in Dutch masker (*t*_(16)_ = 3.31, *p* = 0.002 and *t*_(16)_ = 2.61, *p* = 0.013 for younger listeners in English and Dutch maskers, respectively; *t*_(14)_ = 2.17, *p* = 0.036; *d* = 0.56, 95% *CI* = [3.19×10^−5^, +∞], *t*_(14)_ = 2.55, *p* = 0.061; *d* = 0.66, 95% *CI* = [3.84×10^−5^, +∞] for older listeners in English and Dutch maskers, respectively) (Figure 6B). No significant difference is seen between the slopes across age groups (*t*_(30)_ = 1.36, *p* = 0.091 and *t*_(30)_ = 1.34, *p* = 0.095 for English and Dutch masker, respectively). All tests have been corrected for multiple comparisons across the 6 frequency bands.

**Figure 6.**
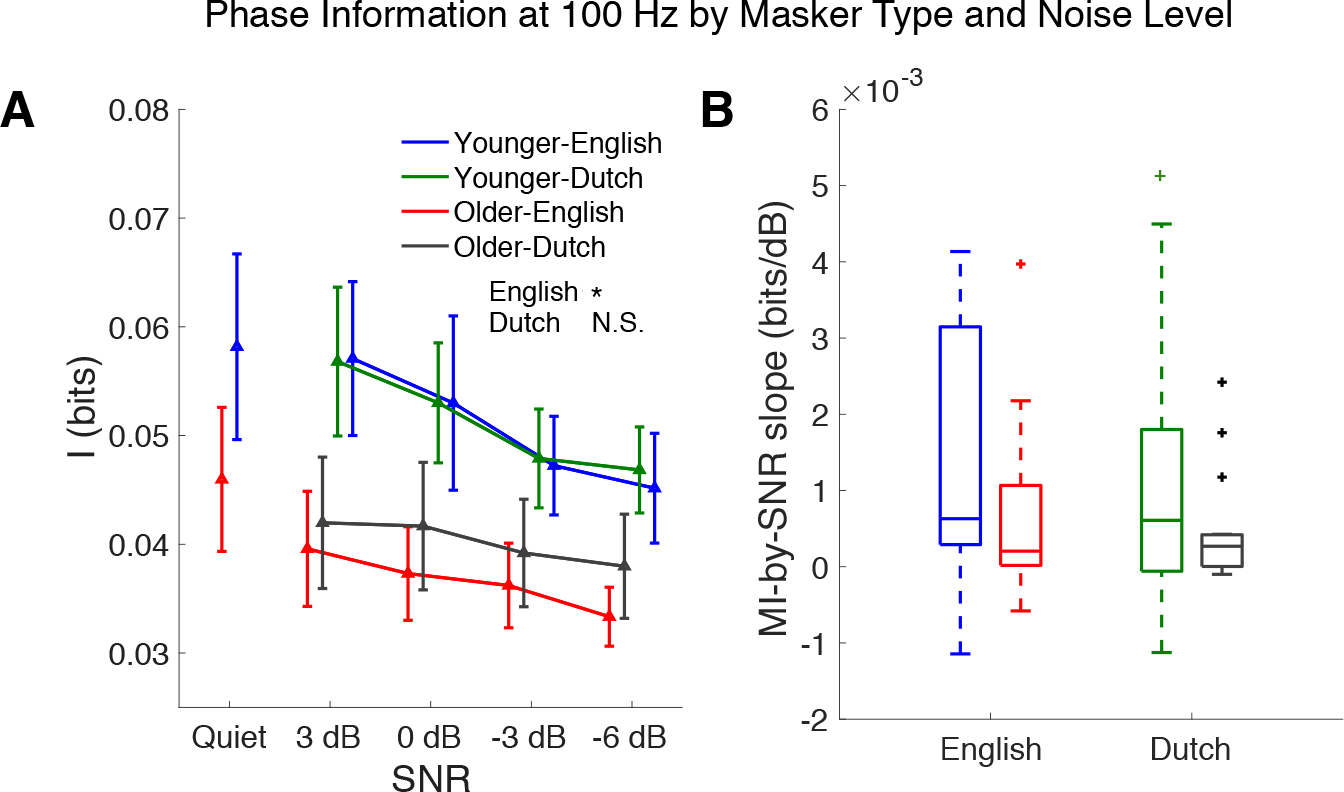
Mutual information between the stimulus and response phase as a function of noise level for each age group and masker condition (masker language). A: Mutual information at the fundamental frequency as a function of noise level. The response in younger listeners conveys noticeably more information than the response in older listeners for the English masker condition, but the difference for Dutch is not significant at 100 Hz. Older listeners show consistently higher mutual information for the Dutch masker than for the English (the younger listeners show no consistent difference), but the difference is not significant at 100 Hz. B: The MI-by-SNR slopes of the previous plots show decreasing trends as SNR worsens, regardless of masker type, for both age groups. Younger listeners show a steeper decrease than older listeners but the difference is not significant at the 100-Hz response. Error bars indicate one SEM. (∗ *p* < 0.05)

#### Phase information in harmonics of 100 Hz

To examine information in the harmonics of 100 Hz, a linear regression line is fitted for mutual information as a function of SNR for each subject under each masker type. One-tailed (younger > older) *t*-tests on the y-intercept (with FDR correction) suggest that for all SNR levels, the aging midbrain contains significantly less information than the younger midbrain in all frequencies from 100 to 600 Hz (Figure 7A). For *p*-values near 0.05 (see Table 2), effect size analysis is further applied. For the English masker condition, the 100 and 200 Hz cases show consistent significance from both tests (*t*_(30)_ = 1.80, *p* = 0.041; *d* = 0.82, 95% *CI* = [0.095, 1.541] and *t*_(30)_ = 1.83, *p* = 0.041; *d* = 1.06, 95% *CI* = [0.317, 1.799]), and similarly for the Dutch masker condition at 300, 400 and 500 Hz, respectively (*t*_(30)_ = 2.12, *p* = 0.042; *d* = 1.39, 95% *CI* = 0.613, 2.159 and *t*_(30)_ = 1.97, *p* = 0.044; *d* = 0.84, 95% *CI* = 0.116, 1.564 and *t*_(30)_ = 2.28, *p* = 0.042; *d* = 1.64, 95% *CI* = [0.838, 2.443]) (see also Figure 7A). The results show significant decreasing slope in both groups and that the decrease with worsening SNR is faster for younger listeners than older listeners.

**Figure 7.**
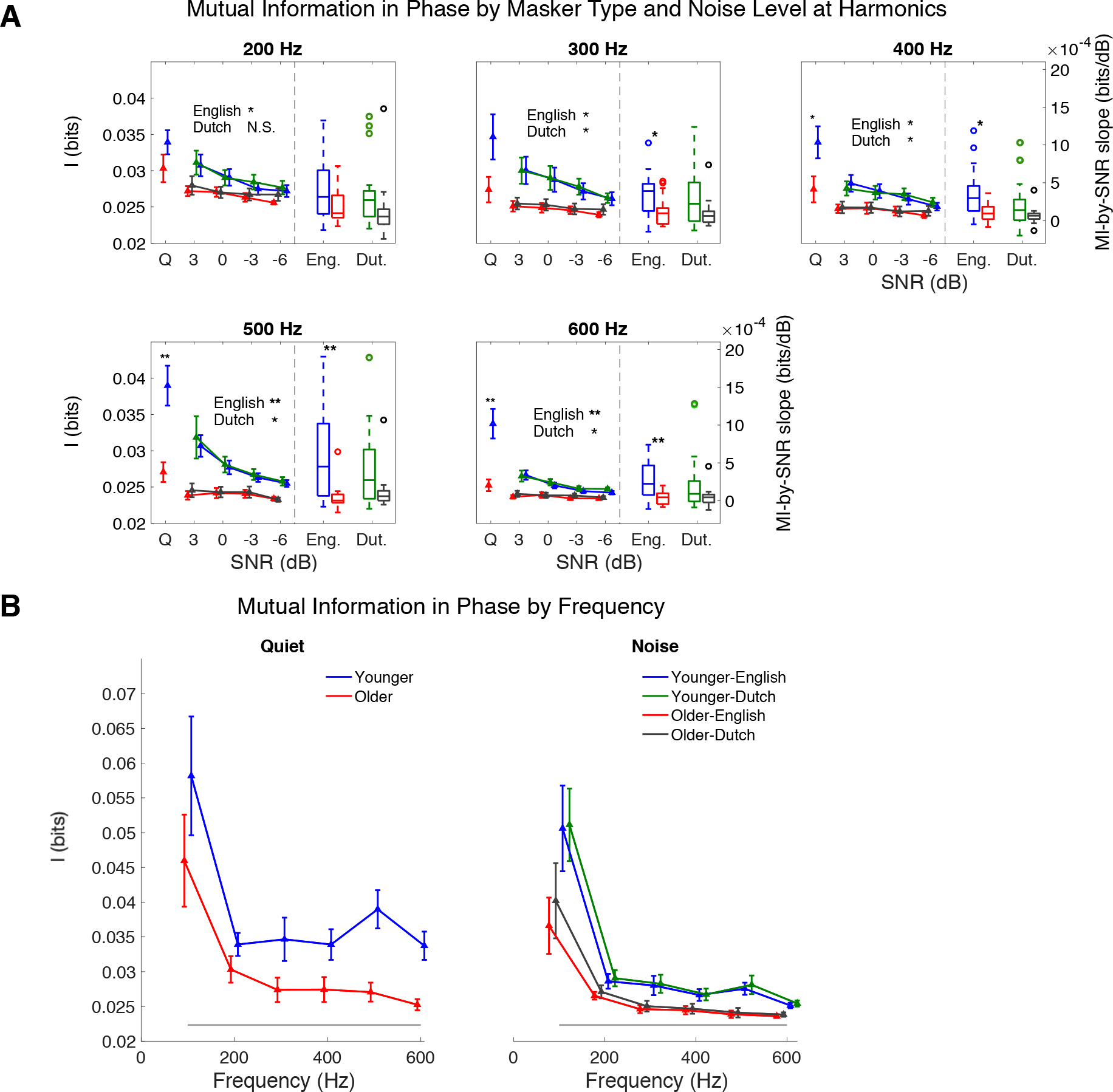
A: Mutual information for phase across frequency bandsfrom 200 Hz to 600 Hz (separate subplot for each band). Within each subplot, as in Figure 3, the left panel shows the mutual information as a function of SNR, separately for age group and masker type; in the right panels, the bar plots depict the linearly fitted decreasing slopes (of the plots shown in the left panel) for the different age groups and masker types. B: Overall, both in quiet (left) and averaged over SNR levels (right), mutual information decreases with increasing frequency (except for a single increase at 500 Hz for younger listeners). For older listeners, the decreasing trend in mutual information levels off at 500 Hz, which is lower than the frequency at which phase information levels off in younger listeners. The lower gray line represents the noise floor. Error bars indicate one SEM. (∗ *p* < 0.05,∗∗ *p* < 0.01)

**Table 2.** Phase information: one-tailed t-test (younger > older) results applied to the fitted yintercepts (3 dB values) and slopes from the linear regression analysis of mutual information (for response phase) as a function of SNR, for each harmonic. p-values are corrected for multiple comparisons by FDR correction. Boldfaced entries indicate the corresponding tests are statistically significant.

| Harmonic<br>(Hz) | Quiet<br>(Y>O) |  | English masker (Y>O) |  |  |  | Dutch masker (Y>O) |  |  |  |
| --- | --- | --- | --- | --- | --- | --- | --- | --- | --- | --- |
|  |  |  | y-intercept |  | slope |  | y-intercept |  | slope |  |
| | $t(30)$ | $p$ | $t(30)$ | $p$ | $t(30)$ | $p$ | $t(30)$ | $p$ | $t(30)$ | $p$ |
| 100 | 1.072 | 0.146 | **1.798** | **0.041** | 1.363 | 0.092 | 1.526 | 0.069 | 1.344 | 0.095 |
| 200 | 1.386 | 0.106 | **1.833** | **0.041** | 1.757 | 0.053 | 1.530 | 0.069 | 1.479 | 0.090 |
| 300 | **1.898** | **0.050** | **2.219** | **0.026** | **2.089** | **0.034** | **2.122** | **0.042** | 1.909 | 0.090 |
| 400 | **2.170** | **0.038** | **2.407** | **0.022** | **2.694** | **0.011** | **1.967** | **0.044** | 1.493 | 0.090 |
| 500 | **3.609** | **0.002** | **3.740** | **0.001** | **3.352** | **0.003** | **2.280** | **0.042** | 1.615 | 0.090 |
| 600 | **3.579** | **0.002** | **3.738** | **0.001** | **3.446** | **0.003** | **2.690** | **0.035** | 1.716 | 0.090 |

#### Phase information frequency limits

As seen in Figure 7B, the stimulus information contained in the response phase decreases with frequency for both age groups. Similar to amplitude analysis, the frequency-decreasing measure used here is phase information of y-intercept at 3dB of the fitted MI-by-SNR regression line. The measure at 600 Hz for older listeners is not statistically distinguishable from the noise floor (*t*_(14)_ = 0.11, *p* = 0.917 by one-sample t-test). For younger listeners, the measure is significantly higher than the noise floor at all frequencies (*t*_(30)_ = 3.74, *p* < 0.001 for English masker; *t*_(30)_ = 2.69, *p* = 0.007 for Dutch masker (younger > older), both at 600 Hz where lowest information is observed), i.e., the information for younger listeners has not yet reached floor by 600 Hz. In contrast, the cutoff frequency for older listeners is 500 Hz: the information measure at 500 Hz is not significantly greater than that at 600 Hz (*t*_(14)_ = 0.74, *p* = 0.235 under English masker; *t*_(14)_ = 1.07, *p* = 0.152 under Dutch masker). Therefore, results suggest a lower frequency limit of 500 Hz for older listeners than beyond 600 Hz for younger listeners.

#### Effect of masker type on phase information

As seen in Figure 6B, older listeners demonstrate a slower fall-off in phase information as a function of SNR when the noise masker is Dutch than for English. Analogous to amplitude analysis, the difference in mutual information between the Dutch and English maskers is calculated for each subject in all SNR levels (for both transition and steady-state regions) to examine phase information benefit from the Dutch masker over the English masker, and a linear model of *I*_*Dutch*_ – *I*_*English*_ ~ *SNR* shows a significantly positive intercept for older listeners in the transition region (*t*_(56)_ = 4.64, *p* < 0.001 with 2 samples omitted) but not in the steady-state region (*t*_(54)_ = 1.77, *p* = 0.083 with 4 samples omitted). Younger listeners, however, do not show significant positive intercept in either transition (*t*_(64)_ = 1.75, *p* = 0.085 with 2 samples omitted) or steadystate region (*t*_(66)_ = −0.64, *p* = 0.522). Samples were omitted from the tests to satisfy the homoscedasticity requirement. For justification, a regression line was fitted as a function of SNR to reduce within-subject variance. Using a one-tailed *t*-test on the y-intercept (effective mutual information benefit at 3 dB SNR) of the regression line against zero, the mutual information benefit from the Dutch masker over the English masker is significantly higher for older listeners in the transition region (*t*_(14)_ = 2.31, *p* = 0.018), but not the steady-state region (*t*_(14)_ = 1.55, *p* = 0.072). No significant benefit is found for younger listeners in either region (*t*_(16)_ = 1.33, *p* = 0.102 and *t*_(16)_ = 0.44, *p* = 0.332 for transition and steady-state region, respectively). The regression slope is not significantly positive or negative for either group (*p* > 0.05 by two-tailed *t*-tests), as seen in the bar plots in the right panels of figure 8C and 8D.

**Figure 8.**
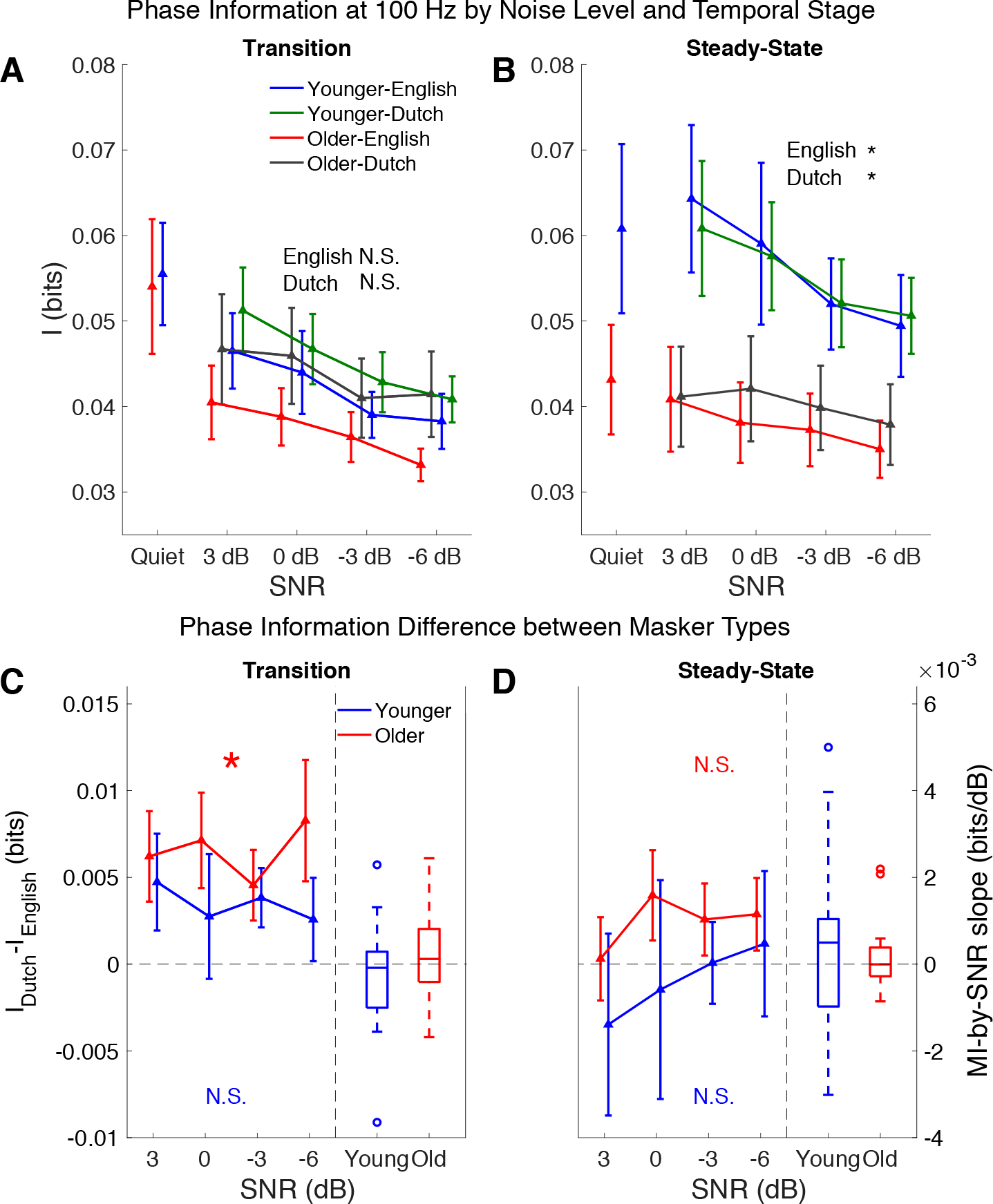
Mutual information of phase response by masker type and response region for younger listeners in blue (English) and green (Dutch) and older in red (English) and gray (Dutch). A and B demonstrate the mutual information as a function of SNR in the transition and steady-stage regions, respectively. In the steady-state region, group differences are significant for both masker types, indicated by asterisks. C and D illustrate the mutual information difference between masker types (denoted *I*_*Dutch*_ – *I*_*English*_) in the transition region and steady-state region, respectively. In each plot, the left panel displays information as a function of SNR, and the right panel displays a bar plot showing the slopes of the linear fits. The y-intercepts (corresponding to the fit at 3 dB SNR) are tested against 0 bits. Older listeners show significant benefit from the Dutch masker over English (denoted by asterisk), but only in the transition region. Error bars in all plots indicate SEM. (∗ *p* < 0.05)

## Discussion

Based on these results from the mutual information analysis of FFR amplitude and phase, we have provided supporting evidence that the neural response of the midbrain of older listeners is not merely less well synchronized than for younger listeners (Anderson et al. 2012; Presacco et al. 2016a, 2016b) but also actually contains less information, in both amplitude and phase. At the fundamental frequency, the informational loss for older listeners was seen only in the presence of a competing talker. In contrast, for higher frequencies, the informational loss for older listeners was seen in both quiet and noisy conditions. Furthermore, the masker type (Dutch vs. English) significantly affects the amount of stimulus information carried in the response at the fundamental frequency in the transition region, for older listeners but not younger. This last finding arises for the first time from this mutual information analysis and demonstrates that mutual information analysis provides access to response properties otherwise hidden by response variability.

### Aging

Aging has different effects on subcortical and cortical auditory stages along the ascending pathway. Here we address its effect on midbrain representations of FFR from an information point of view. First we show a broad-band (100-600 Hz) informational loss associated with aging in both quiet and noisy conditions, which is reflected in both the amplitude and phase of the responses. The informational loss at the fundamental frequency can be attributed to the delayed and weakened responses in the aging midbrain (Anderson et al. 2012; Burkard and Sims 2002; Clinard and Tremblay 2013), which can be linked to age-related loss of inhibition. For example, dorsal cochlear nucleus (DCN) has been shown to represent signal and suppress background noise aided by glycinergic neurotransmitters, and aging rats display decreased glycinergic inhibition in DCN (Caspary et al. 2005, 2006). Another contribution may come from synaptopathy arising from a loss of inner hair cell (IHC) ribbons and degeneration of ganglion cells (Sergeyenko et al. 2013), or from a decline in low-spontaneous-rate nerve fibers as has been seen in aging gerbils (Schmiedt et al. 1996). Together, synaptopathy and loss of inhibition in midbrain may both contribute to less information in midbrain FFR in older listeners.

### Noise level

In these results, the amount of information in FFR (both phase and amplitude) decreases as noise level increases (i.e., SNR decreases) for both younger and older listeners. This result is consistent with previous findings (Presacco et al. 2016a, 2016b) where the amplitude of FFR decreases with worsening noise level. Via linear regression, it is also seen that younger listeners have a more steeply decreasing slope (as a function of noise level) than the older listeners, at both the fundamental frequency and its harmonics. This result may also be due to disrupted synchrony at auditory nerve fibers (Schmiedt et al. 1996) and the synapse (Sergeyenko et al. 2013). A loss of auditory nerve fibers in older listeners may lead to a reduced brainstem response, causing a decrease in information even in the quiet condition, leading to a slower rate of additional decrease with increasing noise level.

### Masker type

In this experiment background masker types included English (meaningful to all listeners) and Dutch (meaningless to all listeners). The results suggest that the informational content of the noise affects information in the midbrain FFR, in both amplitude and phase (in the transition region): older listeners benefit *neurally* from the masker being meaningless over meaningful. It is unexpected that a high-level feature such as language would affect midbrain neural responses, though this has been seen before for younger listeners (Presacco et al. 2016b). One explanation for the language-dependent response difference in the aging midbrain could be top-down modulation from cortical areas. Descending pathways from primary auditory cortex to inferior colliculus (IC) in the midbrain have been reported to mediate learning-induced auditory plasticity (Bajo et al. 2010), and IC neurons’ sensitivity to sound frequency and intensity can be modified by cortical projections (King and Bajo 2013). Since older listeners benefit behaviorally from competing speech being non-meaningful (Pichora-Fuller 2008; Tun et al. 2002), the cortical processing underlying this difference may also project back upstream to the midbrain.

Another explanation for this difference in FFR due to masking language is that the difference might be purely cortical. i.e., purely cortical FFR. Recent studies (Coffey et al. 2016, 2017) have shown that traditional EEG-measured FFR may not be purely subcortical at all. It would be substantially less surprising to see language-specific effects originating from cortex than midbrain, although, even so, these effects from the transition region (15-65 ms) are earlier than might be expected from a language-influenced cortical response.

### High frequency limit

We show that for both amplitude and phase information, responses from older listeners in speech-in-noise conditions contain less information in the higher frequencies, and have lower high frequency limits, than younger listeners. Such deficits might be also associated with lowered temporal precision arising from a loss of auditory nerve fibers and ganglion cells (Schmiedt et al. 1996; Sergeyenko et al. 2013), which affect all frequencies. The same analysis carried out on single sweeps (see Appendix) suggests that the decrease in information at high frequencies may not be due to the average of the two polarities.

### Relation to cortical representation

Even though the stimulus representation at the level of auditory midbrain is weaker for older listeners, whether based on RMS, correlation, or mutual information measures, it is paradoxically amplified at the level of auditory cortex (Brodbeck et al. 2018; Presacco et al. 2016a, 2016b). A negative association between subcortical FFR and cortical responses, as measured with mutual information, has been shown in older listeners in a task of categorical syllable perception (Bidelman et al. 2014). The analogous correlation between cortical speech representation and midbrain response amplitude was not seen, however, for temporal speech processing (Presacco et al, 2016b). Both attention and behavioral inhibition are used to enhance understanding of speech in noise, but the extent to which these high-level cortical processes are altered by auditory periphery deficits is not well known (Presacco et al. 2019). Furthermore, it is unclear where and how the neural representation of speech in older listeners shifts from degraded in midbrain to exaggerated in cortex, but mutual information is a promising tool to address these issues (Bidelman et al. 2014).

### Summary

The approach employed here, using mutual information to analyze the relationship between a speech-in-noise stimulus and the FFR response, can be seen in at least two different lights. At one level it can be viewed as a mathematical measure derived from information theory (Cover and Thomas 1991; Shannon 1948). This places the present analysis on firm mathematical grounds, using concepts and measures from a well-established field of mathematical signal processing. At another level, the analysis can be viewed as an acknowledgement that the relationship between stimulus and response may have strongly non-linear aspects, with mutual information being just one of several available non-linear measures that allow us to move beyond conventional linear analysis methods (e.g. evoked response analysis) and conventional phase coherence methods.

## Appendix

## Results without averaging polarities

Analogously to the case of averaged polarities presented above, even without such polarity averaging, older listeners still demonstrate a slower fall-off in information as a function of SNR when the noise masker is Dutch than for English.

## Information in amplitude of FFR without averaging polarities

For amplitude information, a regression line was fitted as a function of SNR to reduce within-subject variance. Using a one-tailed *t*-test on the y-intercept (effective mutual information benefit at 3 dB SNR) of the regression line against zero, the mutual information in amplitude benefit from the Dutch masker over the English masker is significantly higher for older listeners in the transition region (*t*_(14)_ = 1.80, *p* = 0.046), but not the steady-state region (*t*_(14)_ = 1.61, *p* = 0.065). No significant benefit is found for younger listeners in either region (*t*_(16)_ = 1.04, *p* = 0.156 and (*t*_(14)_ = 0.16, *p* = 0.439 for transition and steady-state region, respectively). The regression slope is not significantly positive or negative for either group (*p* > 0.05 by two-tailed *t*-tests), as seen in the bar plots in the right panels of figure A1C and A1D.

**Figure A1.**
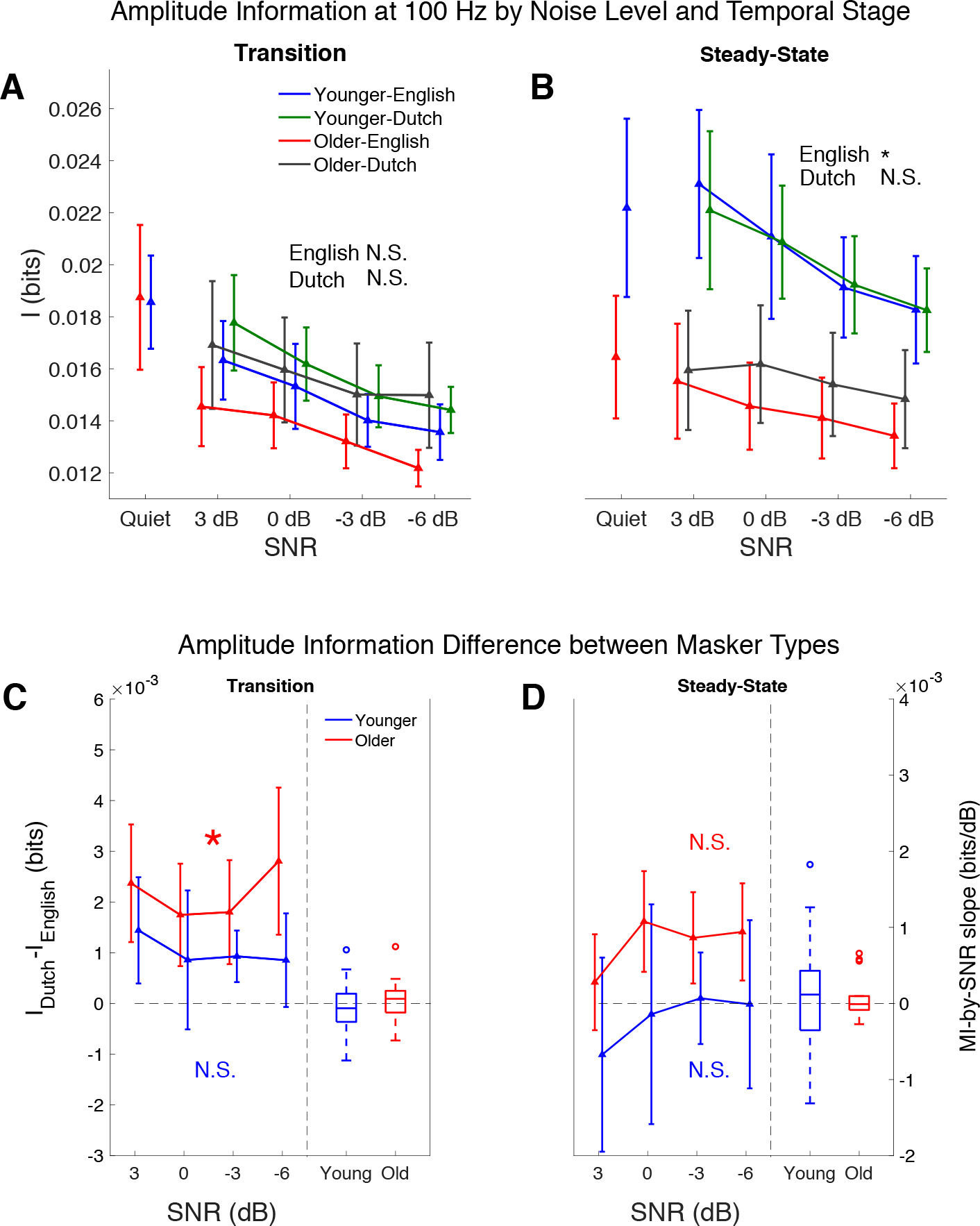
Mutual information of amplitude response by masker type and response region for younger listeners in blue (English) and green (Dutch) and older in red (English) and gray (Dutch). A and B demonstrate the mutual information as a function of SNR in the transition and steady-stage regions, respectively. In the steady-state region, group differences are significant for only the English masker, indicated by asterisks. C and D illustrate the mutual information difference between masker types (denoted *I*_*Dutch*_ – *I*_*English*_) in the transition region and steady-state region, respectively. In each plot, the left panel displays information as a function of SNR, and the right panel displays a bar plot showing the slopes of the linear fits. The y-intercepts (corresponding to the fit at 3 dB SNR) are tested against 0 bits. Older listeners show significant benefit from the Dutch masker over English (denoted by asterisk), but only in the transition region. Error bars in all plots indicate SEM. (∗ *p* < 0.05)

## Information in phase of FFR without averaging polarities

Similarly, for phase information, a regression line was fitted as a function of SNR to reduce within-subject variance. Using a one-tailed *t*-test on the y-intercept (effective mutual information benefit at 3 dB SNR) of the regression line against zero, the mutual information in phase benefit from the Dutch masker over the English masker is significantly higher for older listeners in the transition region (*t*_(14)_ = 1.90, *p* = 0.039), but not the steady-state region (*t*_(16)_ = 1.45, *p* = 0.085). No significant benefit is found for younger listeners in either region (*t*_(16)_ = 1.04, *p* = 0.156 and *t*_(16)_ = 0.25, *p* = 0.401 for transition and steady-state region, respectively). The regression slope is not significantly positive or negative for either group (*p* > 0.05 by two-tailed *t*-tests), as seen in the bar plots in the right panels of figure A2C and A2D.

**Figure A2.**
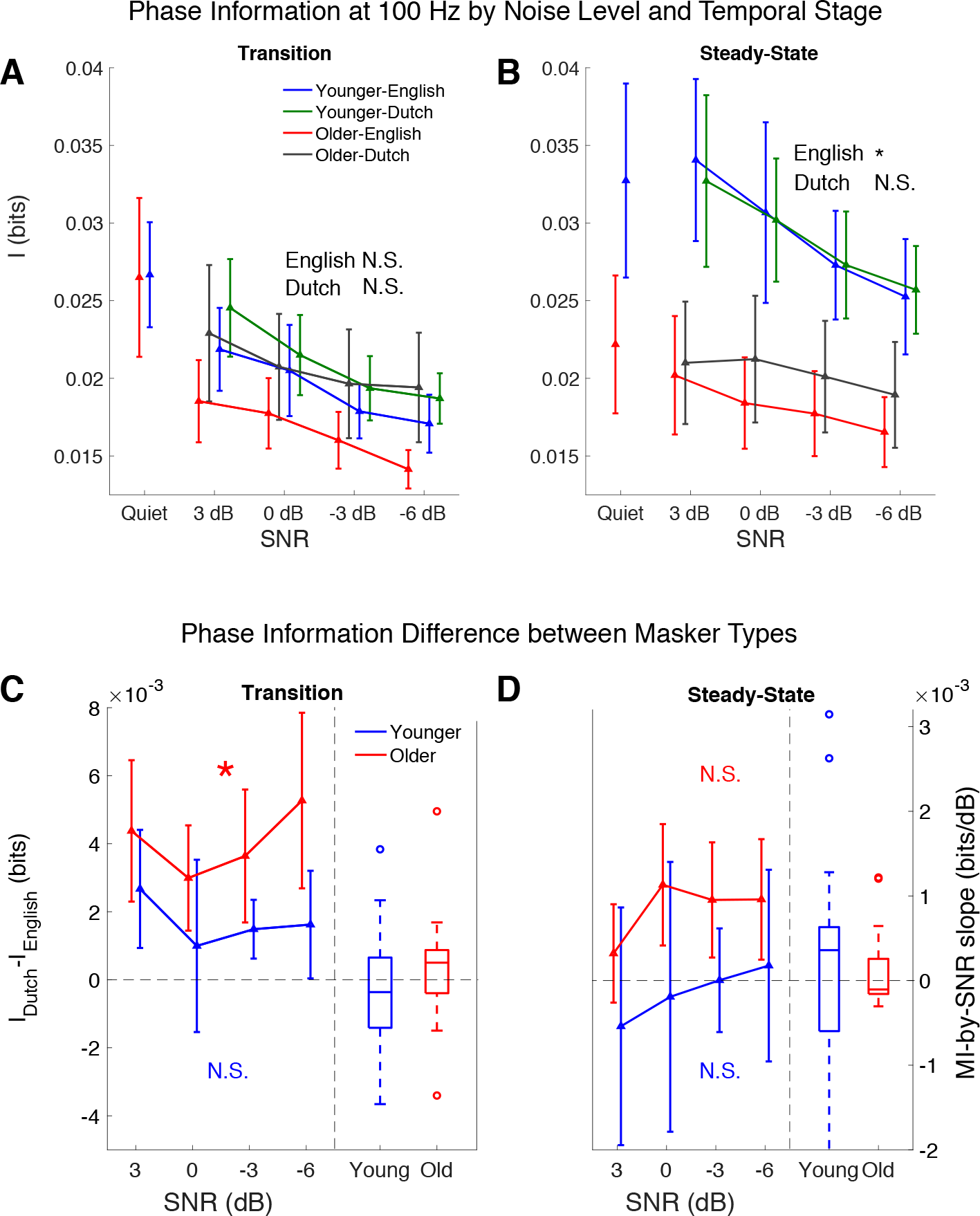
Mutual information of phase response by masker type and response region for younger listeners in blue (English) and green (Dutch) and older in red (English) and gray (Dutch). A and B demonstrate the mutual information as a function of SNR in the transition and steady-stage regions, respectively. In the steady-state region, group differences are significant for only English masker, indicated by asterisks. C and D illustrate the mutual information difference between masker types (denoted *I*_Dutch_ – *I*_*English*_) in the transition region and steady-state region, respectively. In each plot, the left panel displays information as a function of SNR, and the right panel displays a bar plot showing the slopes of the linear fits. The y-intercepts (corresponding to the fit at 3 dB SNR) are tested against 0 bits. Older listeners show significant benefit from the Dutch masker over English (denoted by asterisk), but only in the transition region. Error bars in all plots indicate SEM. (∗ *p* < 0.05)

**Table A1.**
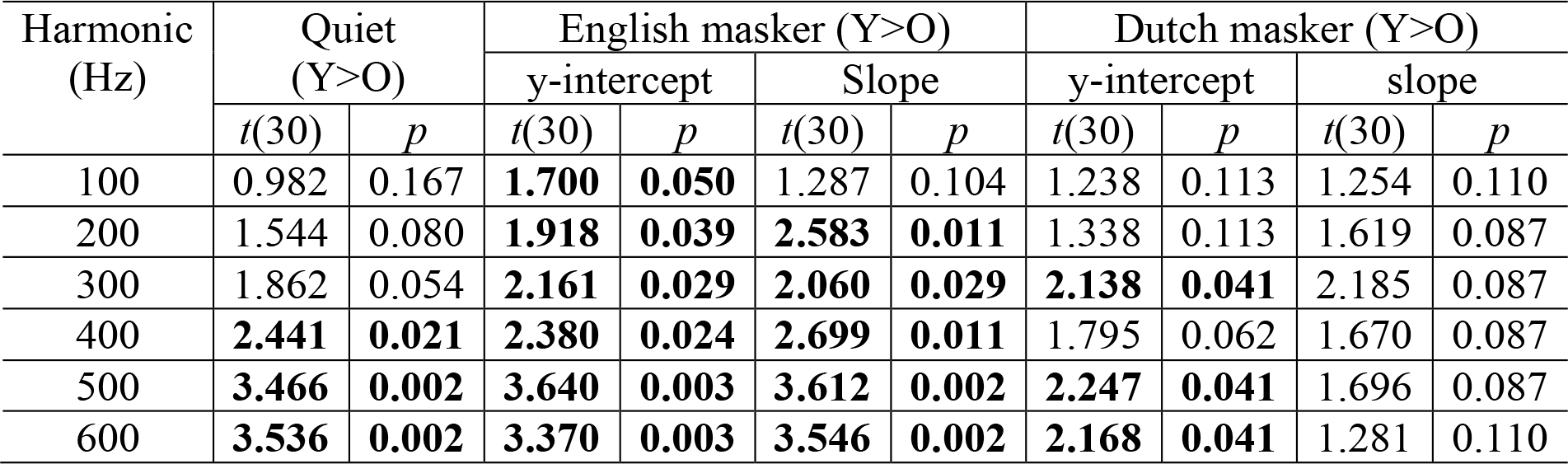
Amplitude information: one-tailed *t*-test (younger > older) results applied to the fitted y-intercepts (3 dB values) and slopes from the linear regression analysis of mutual information (for response amplitude) as a function of SNR, for each harmonic. *p*-values are corrected for multiple comparisons by FDR correction. Boldfaced entries indicate the corresponding tests are statistically significant.

| Harmonic<br>(Hz) | Quiet<br>(Y>O) |  | English masker (Y>O) |  |  |  | Dutch masker (Y>O) |  |  |  |
| --- | --- | --- | --- | --- | --- | --- | --- | --- | --- | --- |
|  |  |  | y-intercept |  | Slope |  | y-intercept |  | slope |  |
| | $t(30)$ | $p$ | $t(30)$ | $p$ | $t(30)$ | $p$ | $t(30)$ | $p$ | $t(30)$ | $p$ |
| 100 | 0.982 | 0.167 | **1.700** | **0.050** | 1.287 | 0.104 | 1.238 | 0.113 | 1.254 | 0.110 |
| 200 | 1.544 | 0.080 | **1.918** | **0.039** | **2.583** | **0.011** | 1.338 | 0.113 | 1.619 | 0.087 |
| 300 | 1.862 | 0.054 | **2.161** | **0.029** | **2.060** | **0.029** | **2.138** | **0.041** | 2.185 | 0.087 |
| 400 | **2.441** | **0.021** | **2.380** | **0.024** | **2.699** | **0.011** | 1.795 | 0.062 | 1.670 | 0.087 |
| 500 | **3.466** | **0.002** | **3.640** | **0.003** | **3.612** | **0.002** | **2.247** | **0.041** | 1.696 | 0.087 |
| 600 | **3.536** | **0.002** | **3.370** | **0.003** | **3.546** | **0.002** | **2.168** | **0.041** | 1.281 | 0.110 |

**Table A2.** Phase information: one-tailed *t*-test (younger > older) results applied to the fitted y-intercepts (3 dB values) and slopes from the linear regression analysis of mutual information (for response phase) as a function of SNR, for each harmonic. *p*-values are corrected for multiple comparisons by FDR correction. Boldfaced entries indicate the corresponding tests are statistically significant.

| Harmonic<br>(Hz) | Quiet<br>(Y>O) |  | English masker (Y>O) |  |  |  | Dutch masker (Y>O) |  |  |  |
| --- | --- | --- | --- | --- | --- | --- | --- | --- | --- | --- |
|  |  |  | y-intercept |  | slope |  | y-intercept |  | slope |  |
| | $t(30)$ | $p$ | $t(30)$ | $p$ | $t(30)$ | $p$ | $t(30)$ | $p$ | $t(30)$ | $p$ |
| 100 | 1.005 | 0.161 | **1.758** | **0.044** | 1.334 | 0.096 | 1.313 | 0.100 | 1.302 | 0.101 |
| 200 | 1.514 | 0.084 | **2.047** | **0.030** | **1.962** | **0.035** | 1.782 | 0.051 | 1.947 | 0.061 |
| 300 | 1.822 | 0.059 | **2.300** | **0.021** | **2.008** | **0.035** | **2.199** | **0.034** | 2.167 | 0.061 |
| 400 | **2.400** | **0.023** | **2.537** | **0.017** | **2.512** | **0.018** | **2.088** | **0.034** | 2.054 | 0.061 |
| 500 | **3.653** | **0.001** | **3.865** | **0.001** | **3.641** | **0.002** | **2.204** | **0.034** | 1.556 | 0.081 |
| 600 | **3.701** | **0.001** | **3.677** | **0.001** | **3.904** | **0.001** | **2.619** | **0.034** | 1.533 | 0.081 |

